# Single-cell transcriptomic profiling of human knee cartilage reveals branching chondrocyte state transitions and extracellular matrix–centered remodeling in osteoarthritis

**DOI:** 10.64898/2026.06.24.734199

**Authors:** Bo Zhu, Xu Han, Yunqi Liang

## Abstract

**Background:** Osteoarthritis cartilage contains heterogeneous chondrocyte states, but molecular programs linked to state transitions within human cartilage remain incompletely resolved using public single-cell data.

**Methods:** A retrospective reanalysis was conducted of a public human knee cartilage single-cell RNA sequencing dataset (GSE255460) including 8 osteoarthritis donors and 3 non-osteoarthritis donors (19 samples). Cells underwent sample-wise quality control and doublet removal, followed by batch-corrected clustering, chondrocyte subclustering with marker-based annotation, and trajectory inference using Slingshot. Regulatory chondrocytes were prioritized for osteoarthritis versus control differential expression, with downstream Gene Ontology/KEGG enrichment (Benjamini–Hochberg false discovery rate <0.05) and protein–protein interaction network hub screening.

**Results:** After quality control, 27,036 cells were retained. Chondrocytes formed multiple transcriptional states with branching-like continuous relationships, and regulatory chondrocytes localized near the main manifold and adjacent to multiple inferred branches, consistent with a transition-adjacent state. In regulatory chondrocytes, osteoarthritis versus control differential expression was enriched for collagen-containing extracellular matrix and extracellular matrix organization, endoplasmic reticulum lumen–associated secretory/proteostasis processes, cell–matrix adhesion (including focal adhesion), and transforming growth factor beta/SMAD-related signaling. Protein–protein interaction analysis of regulatory-chondrocyte differential genes identified five high-connectivity hub genes: COL5A1, COL5A2, COL6A1, COL1A2, and COL3A1.

**Conclusion:** This public-dataset reanalysis supports a transition-adjacent regulatory chondrocyte program in osteoarthritis characterized by coordinated extracellular matrix remodeling with concurrent secretory/proteostasis and adhesion–transforming growth factor beta signatures, nominating collagen-network hubs as candidates for downstream validation.

## Introduction

Osteoarthritis (OA) causes progressive cartilage failure and persistent pain, yet disease-modifying therapies remain elusive, in part because cartilage degeneration reflects coordinated shifts in chondrocyte states and extracellular matrix (ECM) remodeling rather than a uniform chondrocyte phenotype^[1]^. Recent multi-omic and transcriptomic studies support molecular stratification in OA and highlight that ECM-, adhesion-, and stress-response programs vary across tissues and patient subgroups, complicating target prioritization from bulk signals alone^[2,3]^. A major bottleneck is identifying which chondrocyte programs sit near state-transition regions and may coordinate remodeling programs that are potentially tractable for intervention^[4]^.

Single-cell RNA sequencing enables unbiased resolution of cellular heterogeneity and provides a framework to infer continuous, branching-like relationships among cell states within diseased tissues^[4]^. In human OA cartilage, single-cell profiling has delineated multiple molecularly defined chondrocyte populations and suggested potential transitions among proliferative, prehypertrophic, and hypertrophic states, supporting the concept of coordinated state reprogramming during disease progression^[4]^. More recently, integrative analyses combining single-cell and spatial transcriptomics proposed a standardized knee cartilage chondrocyte taxonomy and highlighted several OA-critical populations, but transition-adjacent programs within the broader state space remain incompletely characterized in a way that yields focused, testable candidate networks from public datasets^[4]^.

Here, we conducted a retrospective reanalysis of the public human knee cartilage single-cell RNA sequencing dataset GSE255460, comprising 19 samples from 8 OA donors and 3 non-OA donors^[5]^. After sample-wise quality control and doublet removal, we integrated cells with batch correction, defined chondrocyte states using marker-based annotation aligned to the established knee cartilage framework, and applied trajectory inference to characterize branching-like relationships among chondrocyte states^[5]^. Guided by the inferred state structure, we prioritized a transition-adjacent chondrocyte subpopulation for OA-versus-control comparison and then used functional enrichment and protein–protein interaction network analyses to nominate candidate pathways and hub genes linked to OA-associated ECM remodeling for downstream validation.

## Methods

### Data acquisition and preprocessing

Raw single-cell RNA sequencing (scRNA-seq) count matrices for human knee cartilage were downloaded from the Gene Expression Omnibus (GEO) under accession GSE255460, comprising 19 samples from 8 osteoarthritis (OA) donors (bilateral knees sampled) and 3 non-OA control donors, generated on the 10×Genomics platform (processed feature-barcode matrices)^[5]^. Downstream analyses were conducted in R using Seurat v5.0^[6]^. For each sample, matrices were imported and converted into a Seurat object, and multilayer assays were merged using JoinLayers (Seurat v5 workflow).

To harmonize features across samples, Ensembl gene identifiers were mapped to HGNC gene symbols using org.Hs.eg.db. Features that could not be mapped were removed. Where multiple Ensembl IDs mapped to the same HGNC symbol, duplicated symbols were collapsed by summing raw counts to ensure a unique gene-by-cell count matrix per sample prior to integration. After harmonization and filtering, a total of 22,992 genes were retained for downstream analyses.Clinical trial registration number: Not applicable.

### Quality control and filtering

Quality control (QC) was performed within each sample to minimize cross-sample bias in filtering thresholds. For each cell, QC metrics were computed, including the number of detected genes (nFeature_RNA), total UMI counts (nCount_RNA), mitochondrial transcript fraction (percent.mt, computed using mitochondrial gene prefixes), and ribosomal transcript fraction (percent.ribo). Cells were filtered using fixed thresholds: nFeature_RNA < 200 or > 5,000, percent.mt > 15%, and percent.ribo > 50%.

To mitigate multiplet-driven artifacts, doublets were identified for each sample using scDblFinder (default settings unless otherwise stated), and predicted doublets were removed; only predicted singlets were retained. Following QC and doublet removal, singlets from all samples were merged into a combined object for joint analysis.

To reduce computational burden while preserving per-sample representation, cells were downsampled proportionally by sample, and a minimum retained-cell threshold was applied per sample to avoid the loss of small samples after downsampling. The final integrated dataset used for downstream analyses contained 27,036 cells.

### Normalization, feature selection, dimensionality reduction, and clustering

The merged, downsampled dataset was normalized using Seurat’s standard workflow^[6]^. Highly variable genes (HVGs) were identified, and expression values were scaled prior to dimensionality reduction. Principal component analysis (PCA) was computed using 50 principal components (PCs). Based on the variance-explained profile (elbow-like inflection), the first 40 PCs were retained for downstream analyses.

To correct between-sample batch effects, Harmony was applied using RunHarmony() with sample as the batch covariate^[7]^. Graph-based clustering was performed on the Harmony embedding by constructing a shared nearest-neighbor (SNN) graph (FindNeighbors) and identifying clusters using FindClusters with resolution = 0.5 (Louvain/Leiden depending on Seurat defaults and installed backends). Clusters were visualized using UMAP (RunUMAP) with default parameters. Because clustering and UMAP can be sensitive to tuning parameters, the low-dimensional structure was interpreted as a visualization and clustering aid rather than a direct quantitative model of biological distances^[8]^.

### Cell type annotation and chondrocyte subclustering

Cell type annotation was first performed using SingleR v2.8.0^[9]^ with the Human Primary Cell Atlas as the reference, applied at the cluster level. Automated labels were then reviewed using canonical marker genes. Chondrocytes were subsetted and re-clustered to define chondrocyte subpopulations; subpopulation naming was aligned to the established knee cartilage chondrocyte framework reported for this dataset^[5]^.

### Differential expression analysis and functional enrichment

Marker genes for global clusters and chondrocyte subpopulations were identified using Seurat’s FindMarkers function (default statistical test unless otherwise specified). For primary downstream analyses, regulatory chondrocytes (RegC; CHI3L1/CHI3L2-high) were prioritized based on their trajectory-adjacent positioning (see below). Within RegC, differential expression testing was performed comparing OA vs control, yielding RegC-specific OA-associated differentially expressed genes (DEGs).

Functional enrichment analysis was conducted using clusterProfiler. Over-representation analyses were performed for Gene Ontology (GO; BP/CC/MF) and KEGG pathways using the RegC DEG list as input. Multiple testing correction was performed using the Benjamini–Hochberg method, and statistical significance was defined as FDR (q value) < 0.05.

### Trajectory inference

To characterize branching-like state relationships among chondrocyte subpopulations, pseudotime trajectory inference was performed using Slingshot v2.14.0 on the chondrocyte subset. Slingshot was run using the low-dimensional embedding (Harmony/UMAP or PCA-derived reduced dimensions, consistent with the analysis pipeline) and cluster labels as inputs to infer lineage structure and pseudotime ordering. Branch-associated expression patterns were examined to support interpretation of state adjacency and branching organization. Consistent with current best practices, pseudotime was used to describe state proximity and branching structure rather than to assert definitive biological start/end directionality^[4]^.

### Protein–protein interaction network analysis and hub gene identification

RegC OA-associated DEGs were uploaded to STRING^[10]^ v11.0 (https://string-db.org/) to retrieve known and predicted protein–protein interactions. The resulting interaction network was imported into Cytoscape v3.10.1 for visualization and topological analysis^[11]^. Hub genes were prioritized using the cytoHubba plugin, applying the Degree algorithm to rank nodes by connectivity; the top 5 highest-degree genes were reported as candidate hub genes.

### Reproducibility and reporting

All analyses were performed in R with package versions recorded (e.g., via sessionInfo()), and plotting/code were executed under fixed random seeds where applicable to improve reproducibility. Key intermediate objects (post-QC Seurat objects, Harmony embeddings, cluster labels, DEG tables, enrichment results, and network edge lists) were retained to enable end-to-end reruns of the pipeline and independent verification.

## Results

### Quality control and data retention

To assess data quality and define the analyzable cell set, we first evaluated sequencing depth–feature relationships and per-sample quality-control metrics (Fig. 1–3), following standard scRNA-seq analysis practice for preprocessing and downstream interpretability. After gene identifier harmonization and filtering, a gene-by-cell expression matrix containing 22,992 genes was obtained. After sample-wise quality control, doublet removal, and proportional downsampling to preserve sample representation, 27,036 cells were retained for downstream analyses. Across the retained cells, sequencing depth showed an increasing trend with detected gene features, with a dense low-value region and multiple high-value outliers (Fig. 1). Per-sample distributions of nFeature_RNA and nCount_RNA varied, with most cells concentrated in mid-range values and a subset of high-count outliers (Fig. 3).

**Figure 1.**
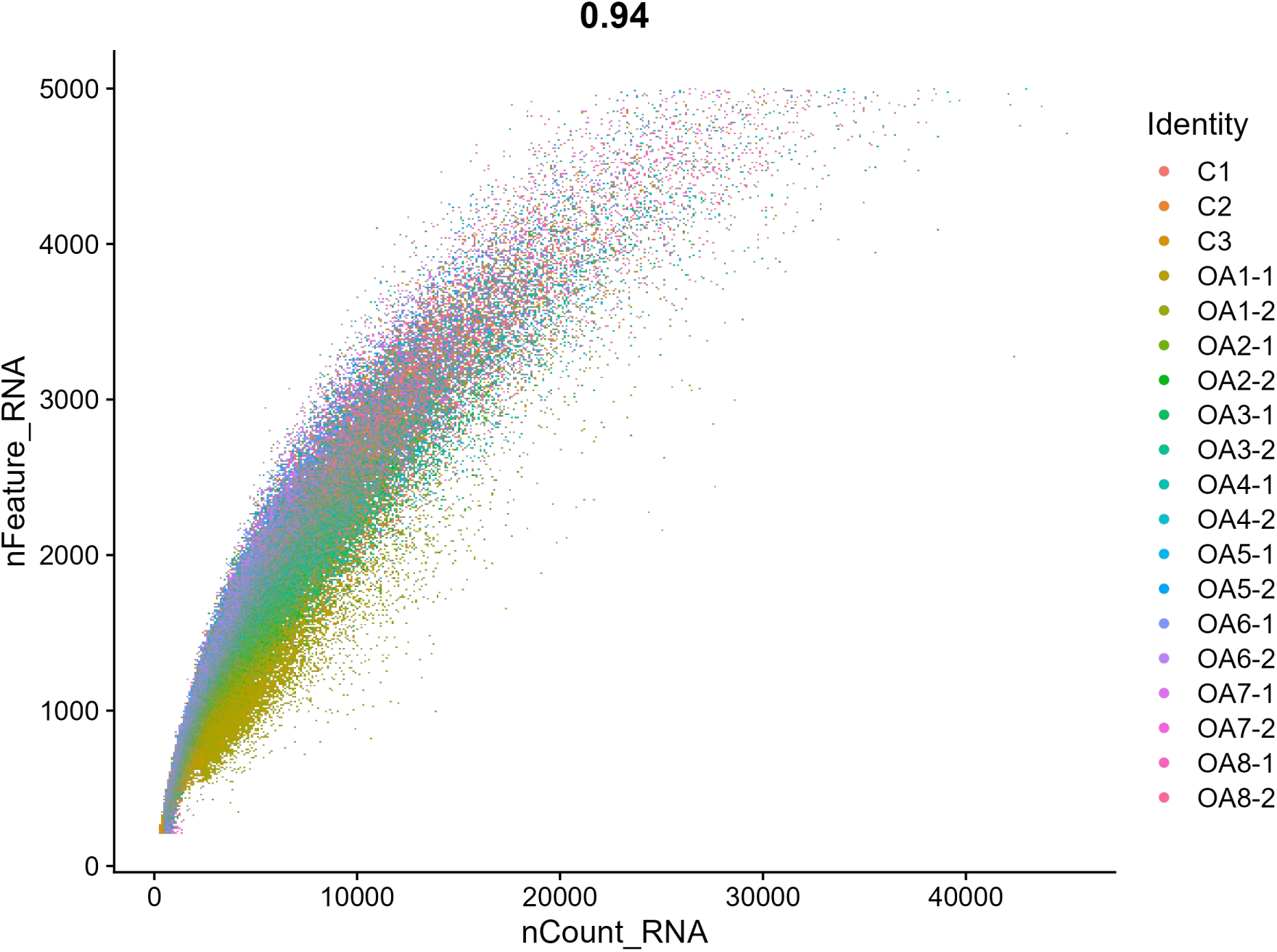
Sequencing depth versus detected gene features after quality control. Scatter plot showing the relationship between sequencing depth and detected gene features, with a dense low-value region and multiple high-value outliers.

**Figure 2.**
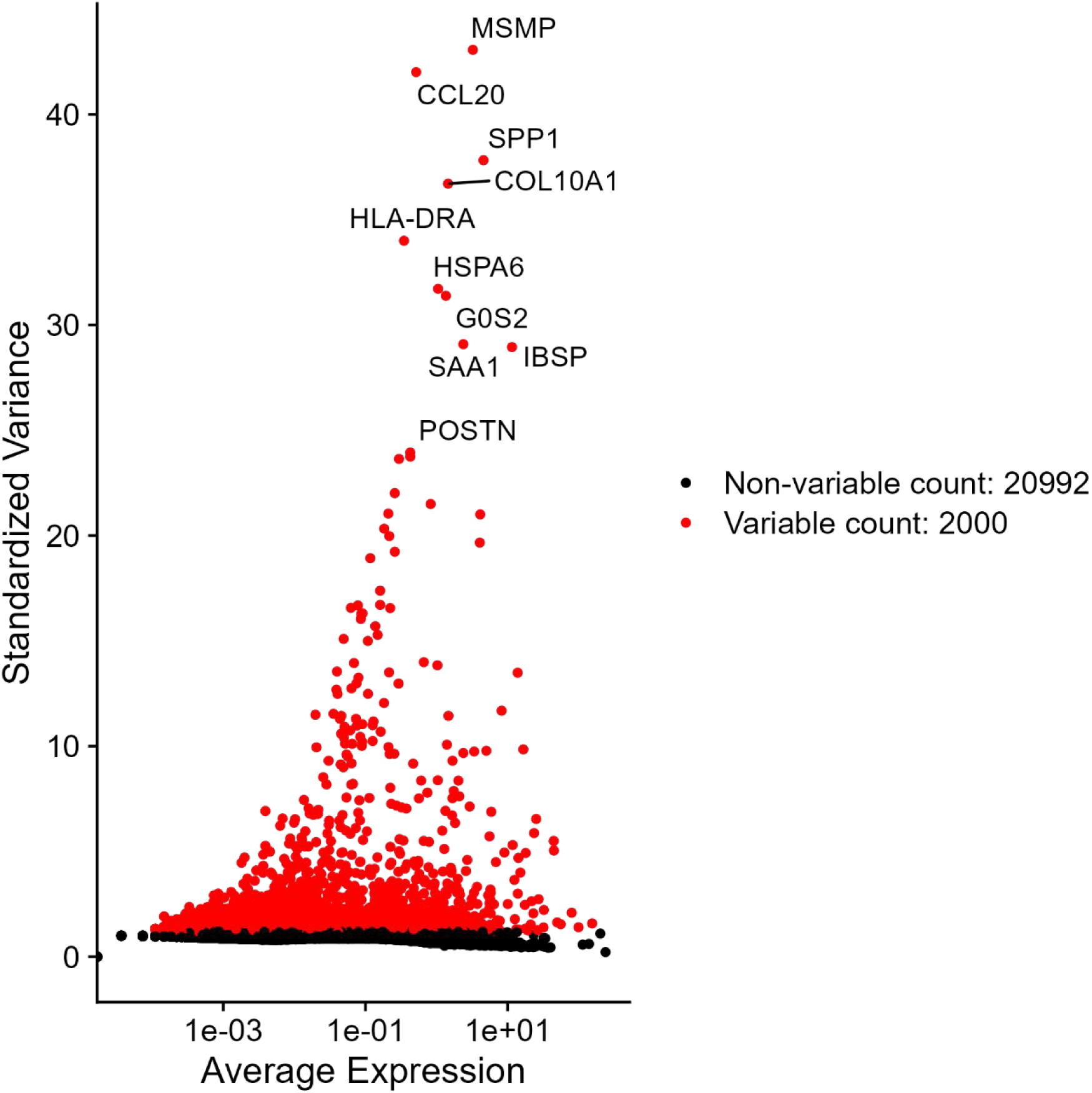
Highly variable gene selection / gene expression overview. Plot showing the selected variable genes used for downstream analyses (as displayed in the figure).

**Figure 3.**
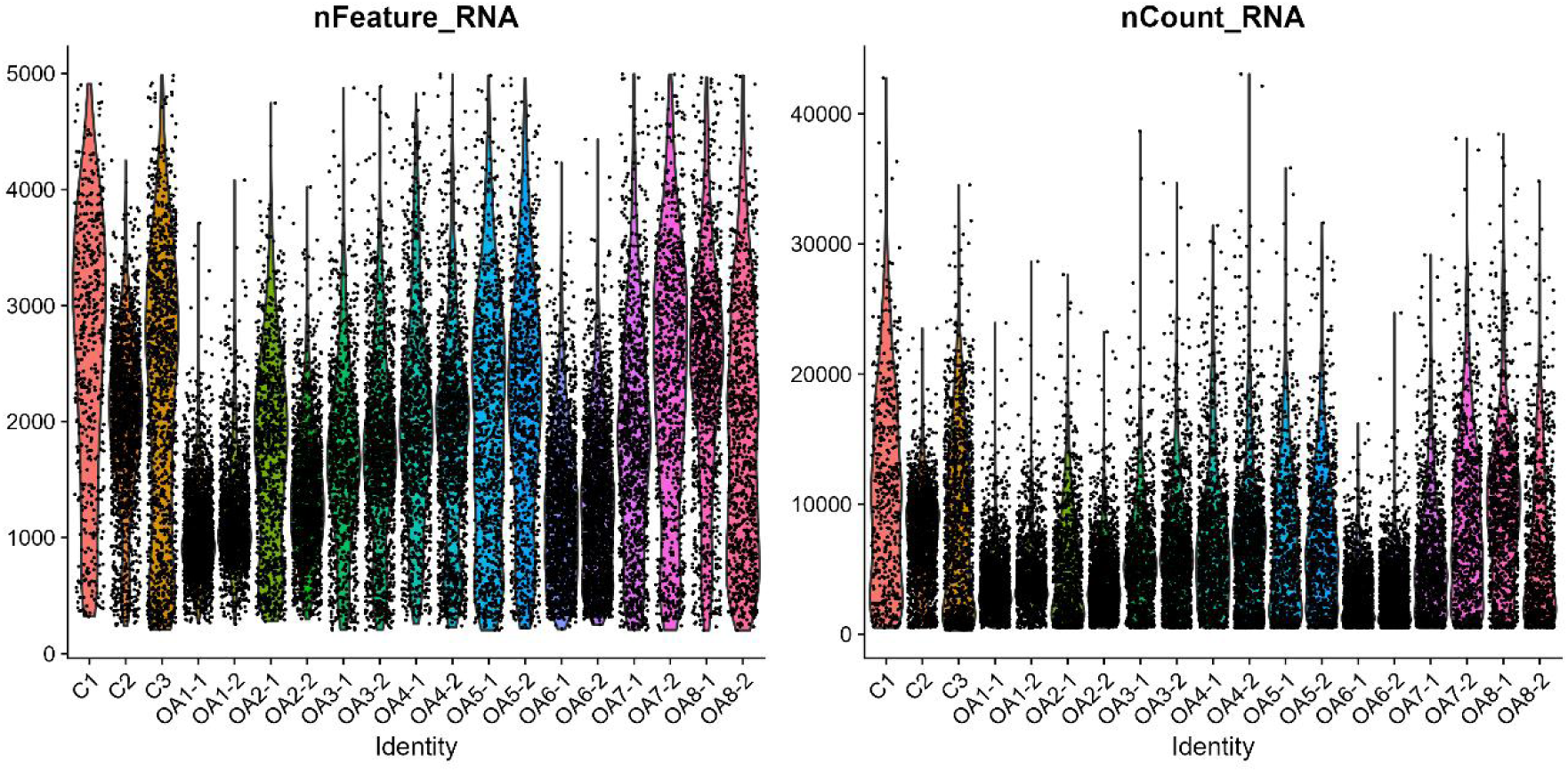
Per-sample distributions of sequencing depth and gene features. Violin/strip plots showing distributions of nFeature_RNA and nCount_RNA across samples after quality control.

### Principal component structure and selection of informative dimensions

To reduce dimensionality for clustering and visualization, we performed principal component analysis (PCA) and selected informative components based on variance structure (Fig. 4–8), consistent with best-practice workflows that use variance-explained patterns to guide component retention. PCA was computed using 50 principal components, and 40 components were retained for downstream analyses based on the variance-explained profile and an elbow-like inflection (Fig. 4–5). Projection on PC1 and PC2 summarized the global structure of chondrocyte transcriptomes in low-dimensional space (Fig. 6). Heatmaps and gene-loading summaries for the top principal components highlighted component-associated genes, including COL1A1 (PC1), TYROBP (PC2), SPARC (PC3), and COMP (PC4) (Fig. 7–8).

**Figure 4.**
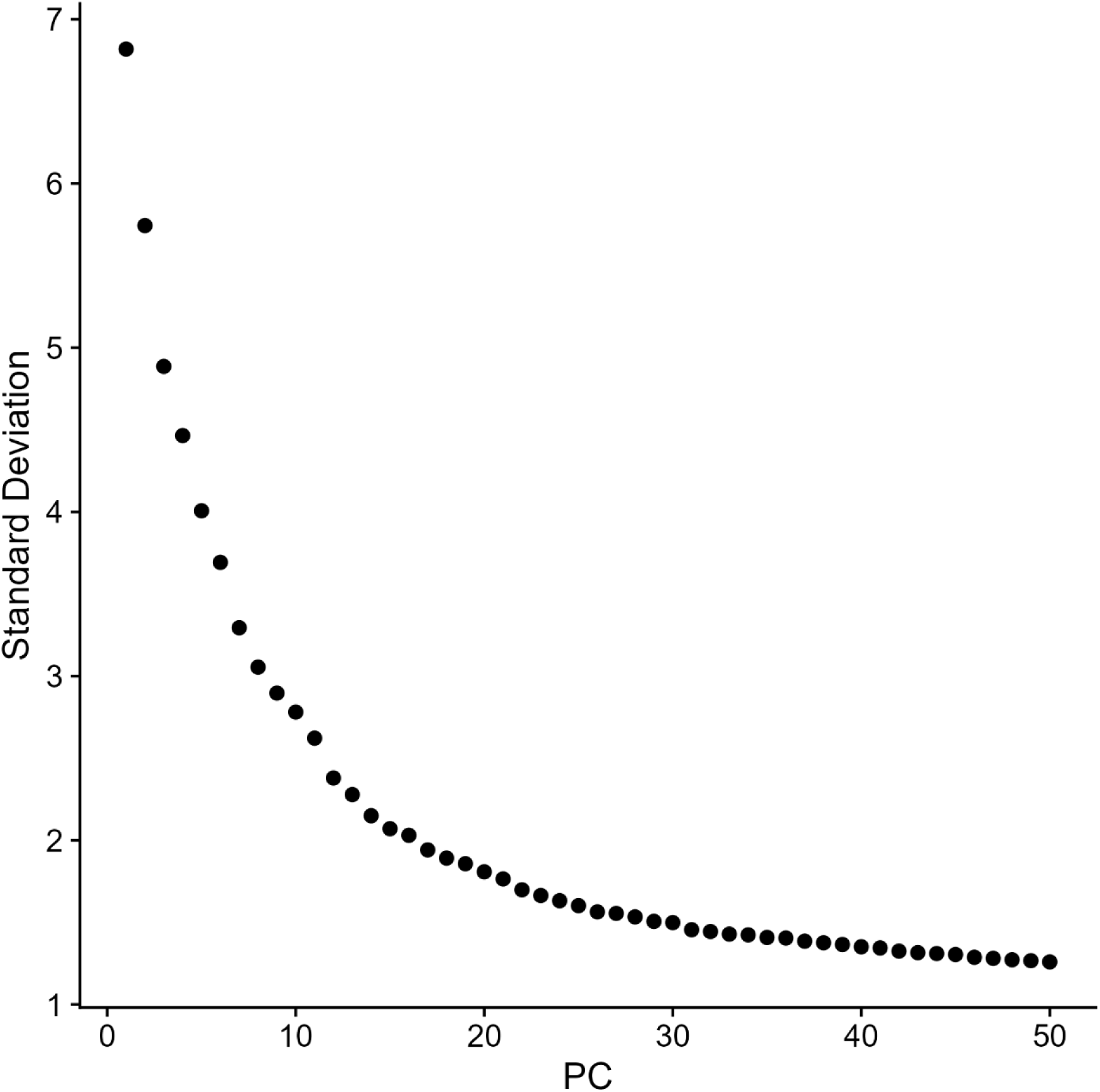
Variance-explained profile across principal components. Elbow/variance plot summarizing variance explained across computed principal components.

**Figure 5.**
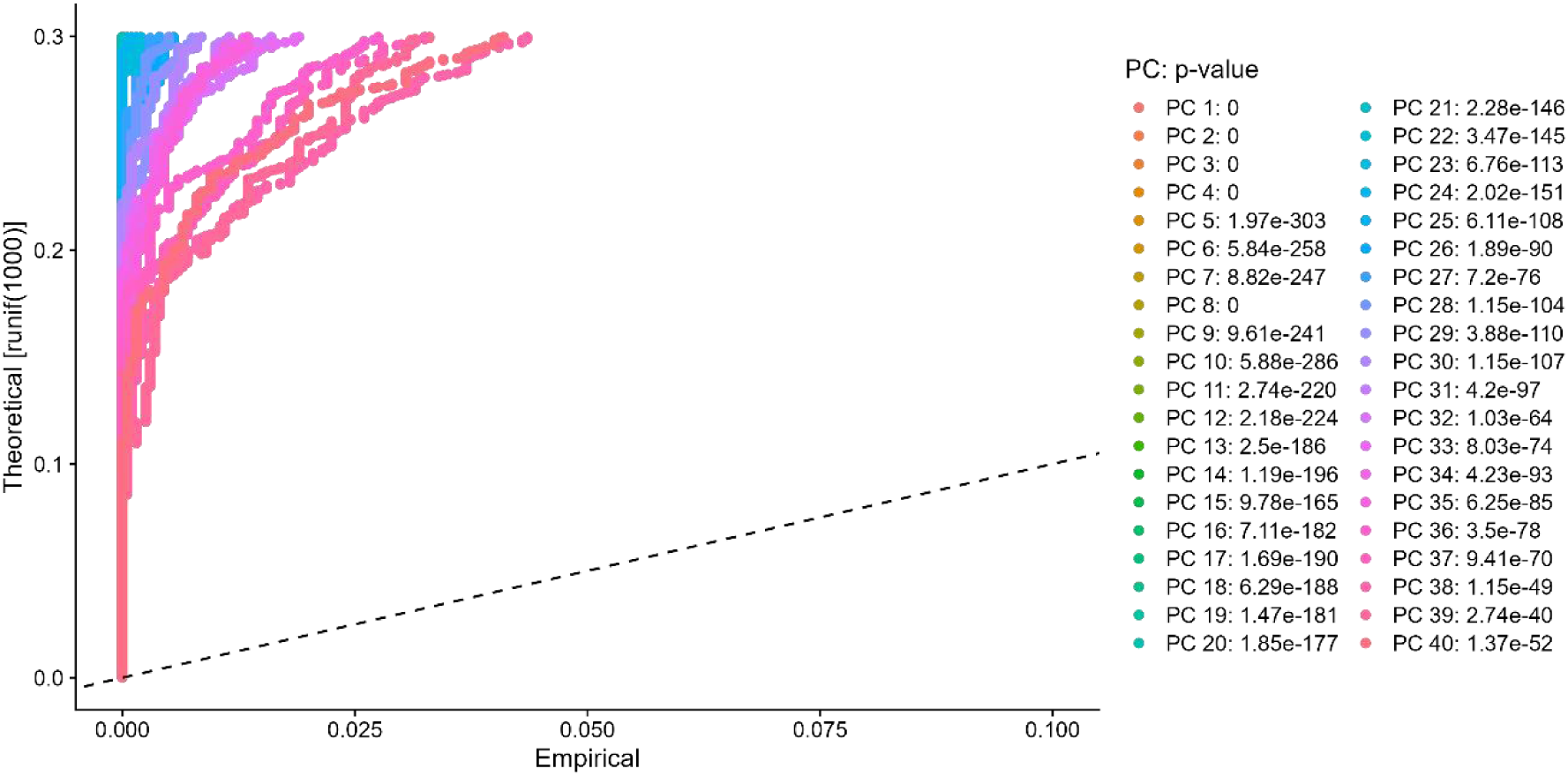
Principal components retained for downstream analyses. Summary plot indicating retention of 40 principal components based on the variance-explained inflection.

**Figure 6.**
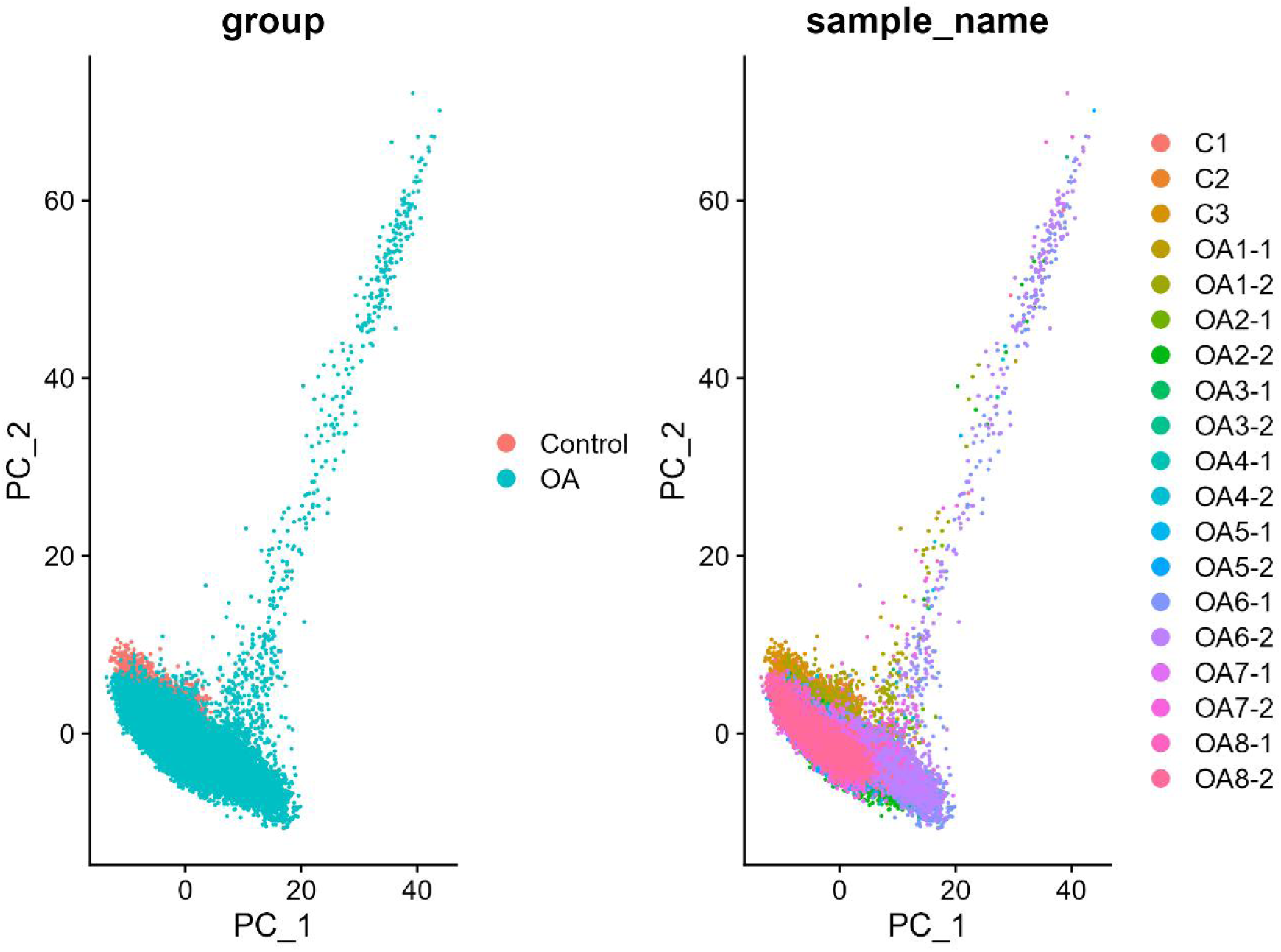
PCA projection of chondrocyte transcriptomes. Scatter plot of cells projected onto PC1 and PC2.

**Figure 7.**
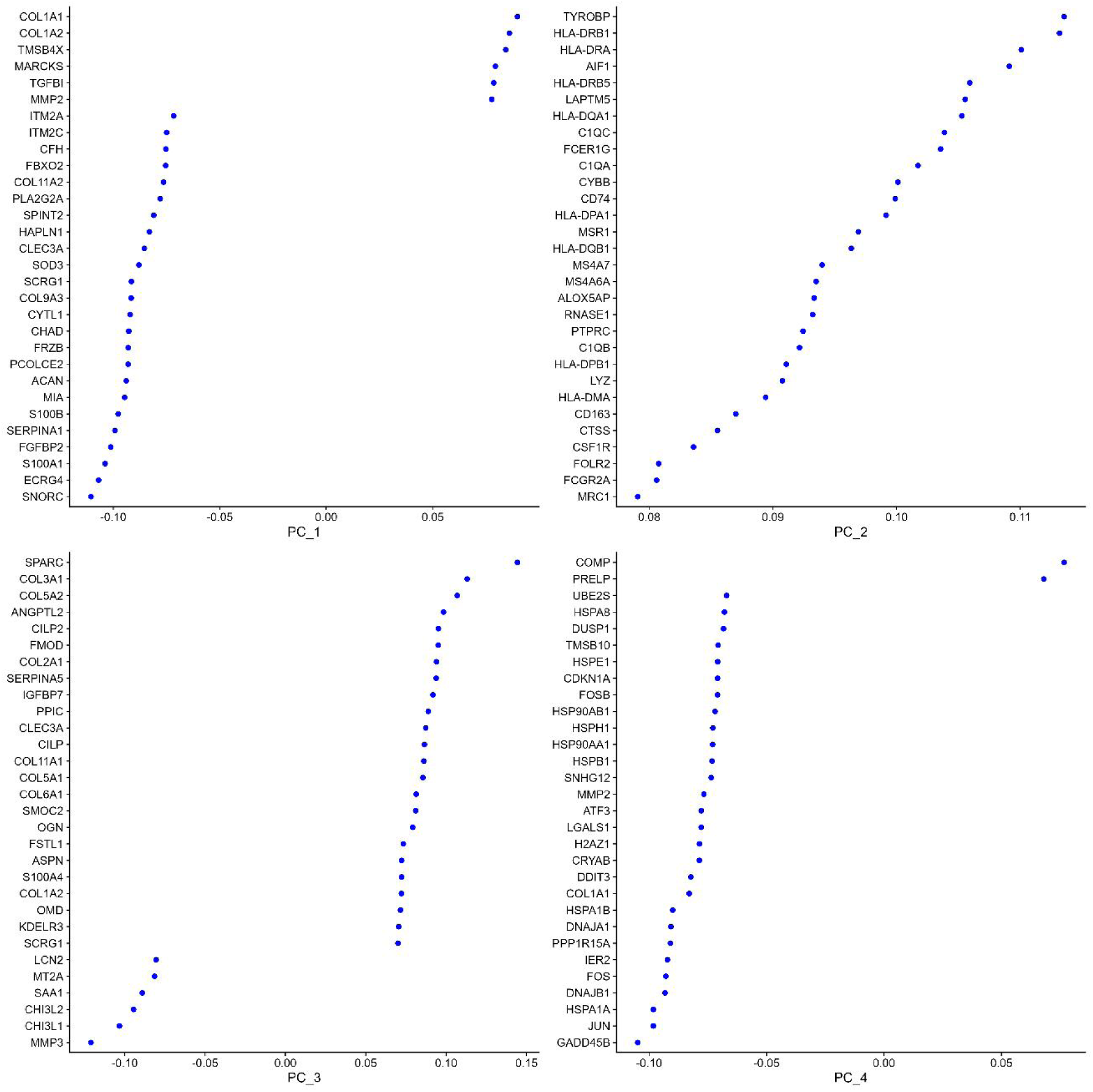
Gene-loading summary for the top principal components. Curves/plots summarizing genes contributing to PC1–PC4.

**Figure 8.**
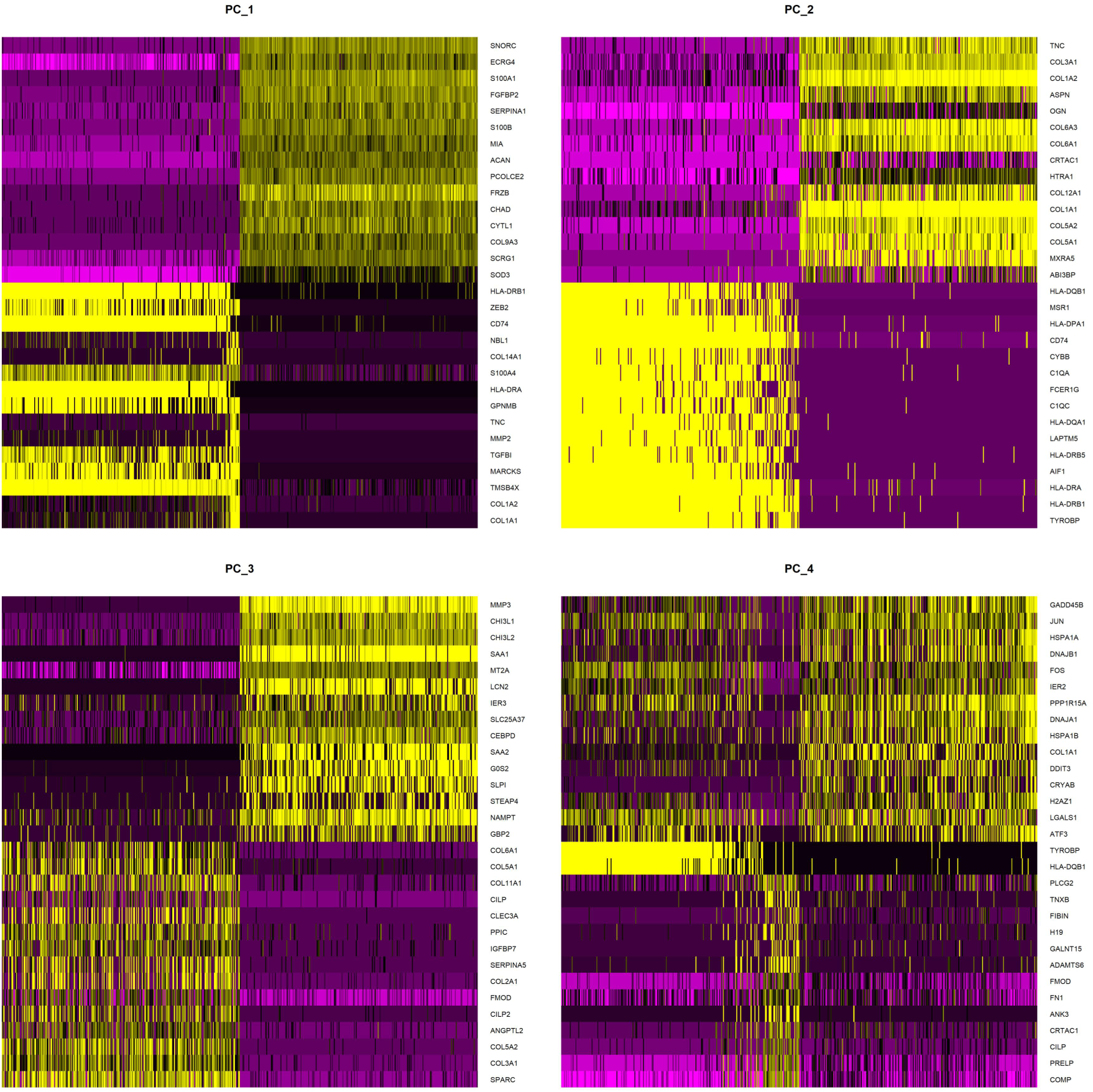
Heatmap of genes associated with the top principal components. Heatmap showing expression patterns of genes contributing to PC1–PC4, including component-associated genes (COL1A1, TYROBP, SPARC, COMP).

### UMAP clustering identifies transcriptionally distinct cell clusters and marker genes

To construct a global single-cell atlas of knee cartilage, we performed batch-corrected integration followed by graph-based clustering and UMAP visualization (Fig. 9–10), aligning with commonly used atlas construction workflows. UMAP embedding resolved 11 transcriptional clusters (Fig. 9). Cluster distributions differed between osteoarthritis (OA) and non-OA control samples, with several clusters present in both groups (Fig. 9). Using an adjusted significance threshold of FDR(q)<0.05, we identified 3,225 cluster marker genes and visualized their expression patterns across clusters (Fig. 10).

**Figure 9.**
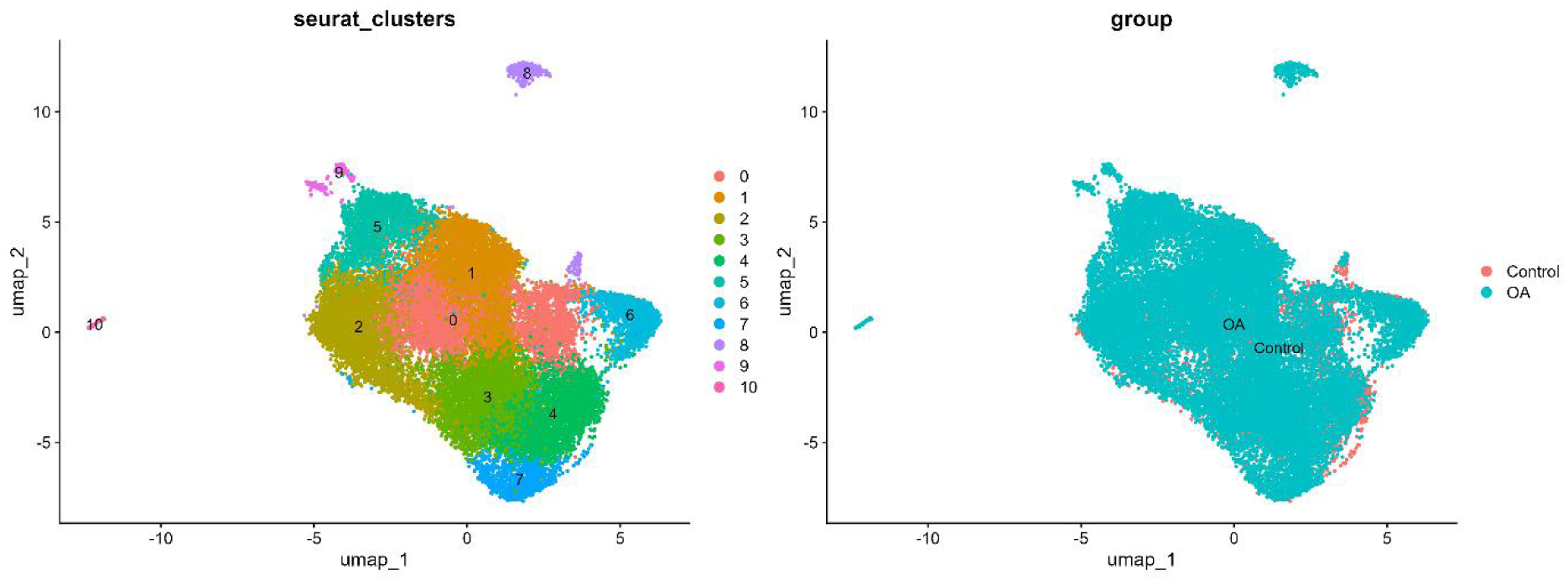
UMAP embedding and cluster distribution across OA and control samples. UMAP visualization showing 11 clusters and their distribution across OA and control groups.

**Figure 10.**
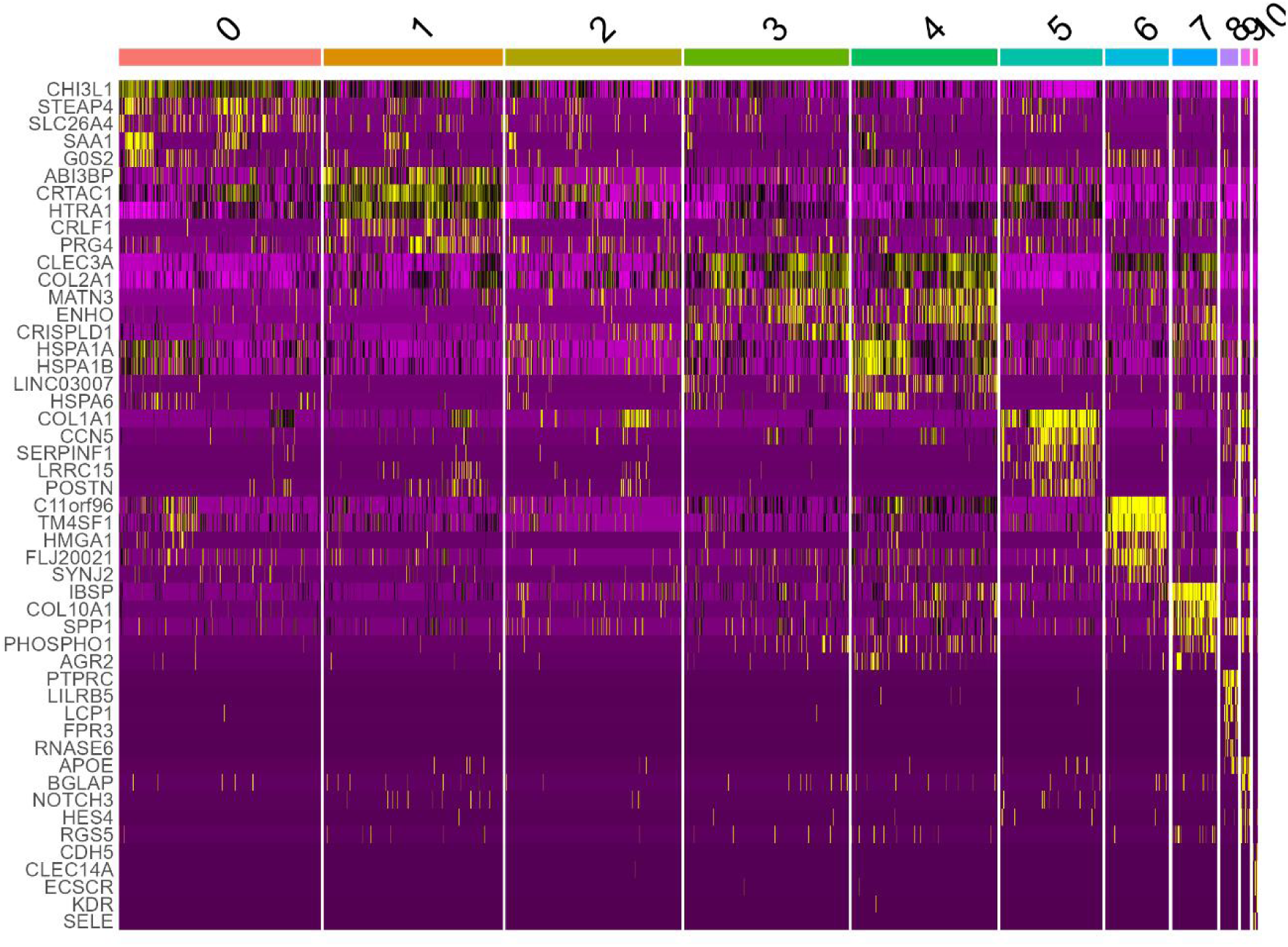
Heatmap of cluster marker genes (FDR(q)<0.05). Heatmap showing expression of 3,225 marker genes across the 11 clusters.

### Cell-type annotation and trajectory inference delineate chondrocyte subpopulations and branching structure

To assign biological identities to clusters and characterize state relationships within chondrocytes, we performed automated annotation with SingleR followed by marker-based refinement and trajectory inference (Fig. 11–12). Marker-based annotation defined multiple chondrocyte subpopulations, including effector chondrocytes (EC; CHRDL2, FRZB, CYTL1), fibrocartilage chondrocytes (FC; MMP2, COL1A1, COL1A2), hypertrophic chondrocytes (HTC; SPP1, IBSP, COL10A1), pre-inflammatory chondrocytes (preInfC; IFI16, IFI27), inflammatory chondrocytes (InfC; CXCL8, CD74, GPR183), progenitor-like chondrocytes (ProC; C11orf96, BMP2, HMGA1), and regulatory chondrocytes (RegC; CHI3L1, CHI3L2) (Fig. 11A, Fig. 12). Slingshot trajectory inference identified 5 major branching lineages within the chondrocyte manifold (Fig. 11B). RegC localized near the main trunk region and adjacent to multiple inferred branches (Fig. 11B). This overall pattern of continuous state structure with branching-like relationships is consistent with prior OA cartilage single-cell work reporting chondrocyte state diversity and potential state transitions in human OA cartilage.

**Figure 11.**
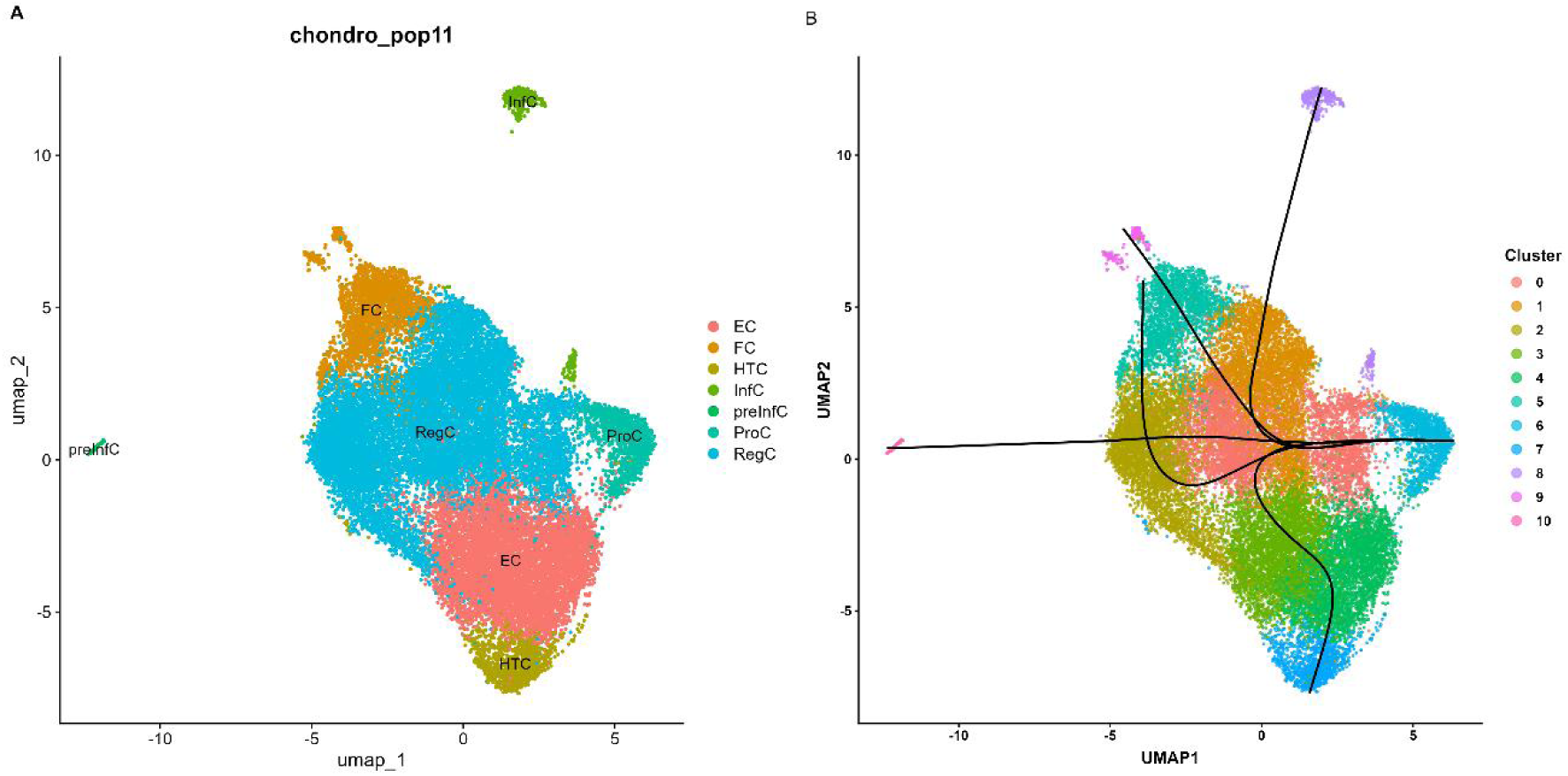
Chondrocyte subpopulation distribution and Slingshot trajectories. (A) UMAP showing annotated chondrocyte subpopulations. (B) Slingshot pseudotime trajectories showing 5 major branching lineages; RegC localized near the trunk and adjacent to multiple branches.

**Figure 12.**
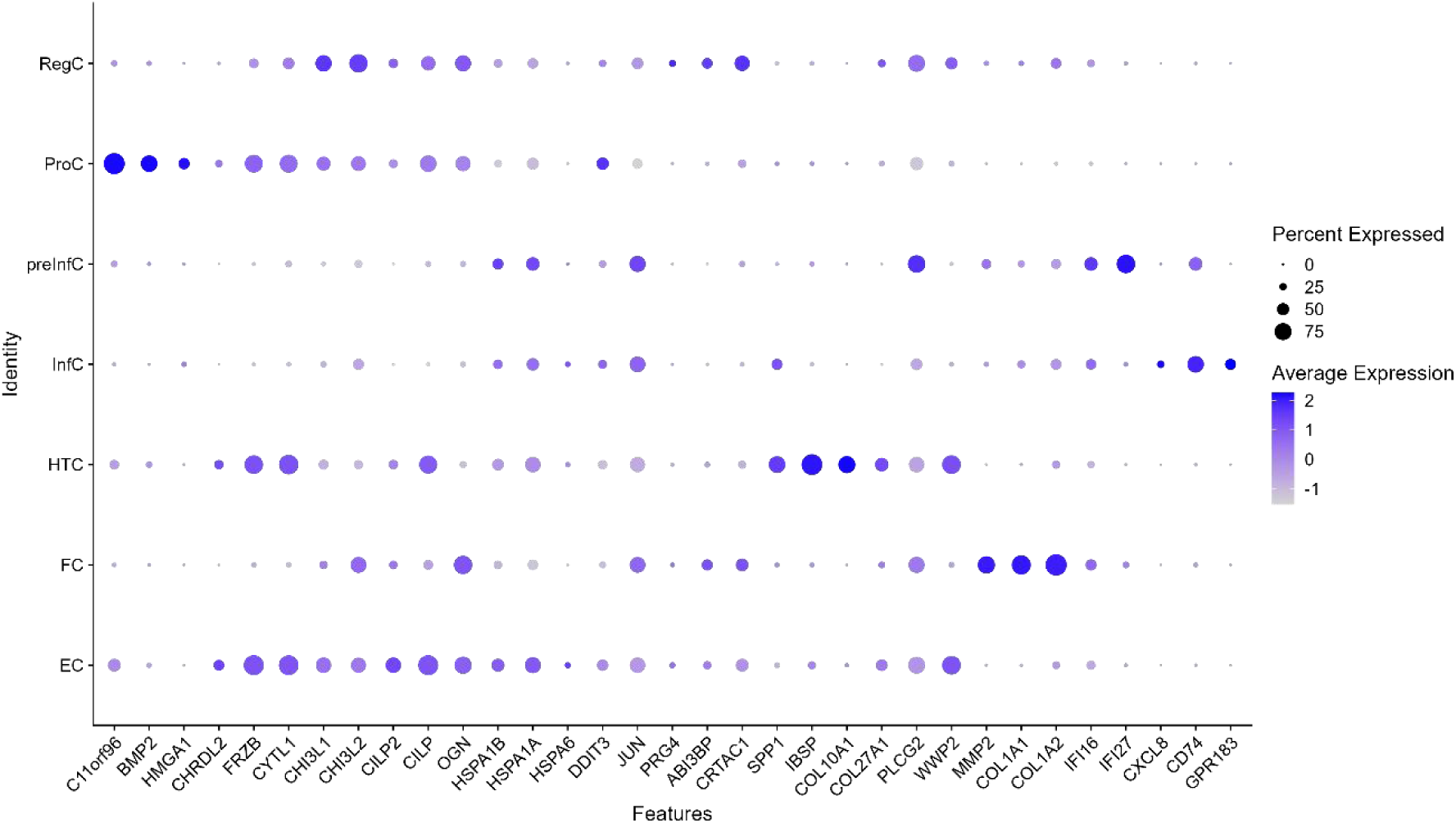
Bubble plot of subpopulation marker genes. Bubble plot showing expression patterns of marker genes across annotated chondrocyte subpopulations.

### Functional enrichment of OA-associated differential genes in transition-adjacent regulatory chondrocytes

To characterize OA-associated transcriptional changes within the transition-adjacent state, we performed differential expression analysis within RegC (OA vs control) followed by functional enrichment (Fig. 13–14). RegC differential genes were enriched for collagen-containing extracellular matrix and extracellular matrix organization terms, endoplasmic reticulum lumen–associated secretory/proteostasis processes, cell–matrix adhesion (including focal adhesion-related terms), and transforming growth factor beta/SMAD-related signaling (Fig. 13–14). Multiple-testing correction was applied using Benjamini–Hochberg, with FDR(q)<0.05 as the significance threshold. The prominence of OA-associated signals in specific chondrocyte populations is consistent with integrative knee cartilage mapping studies that emphasize population-resolved OA-critical programs rather than bulk-average effects.

**Figure 13.**
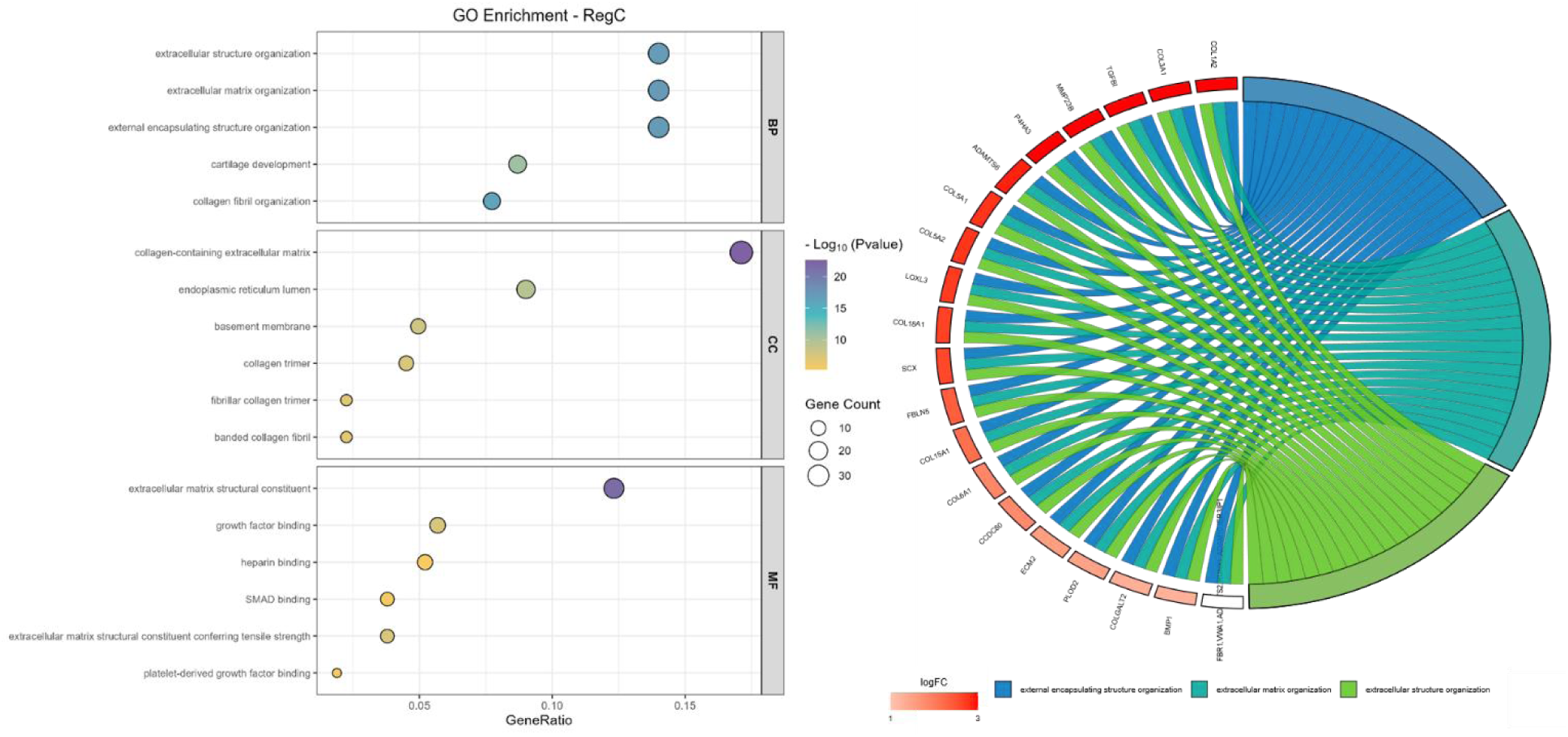
Gene Ontology enrichment of RegC OA-versus-control differential genes (FDR(q)<0.05). GO over-representation analysis results for RegC differential genes.

**Figure 14.**
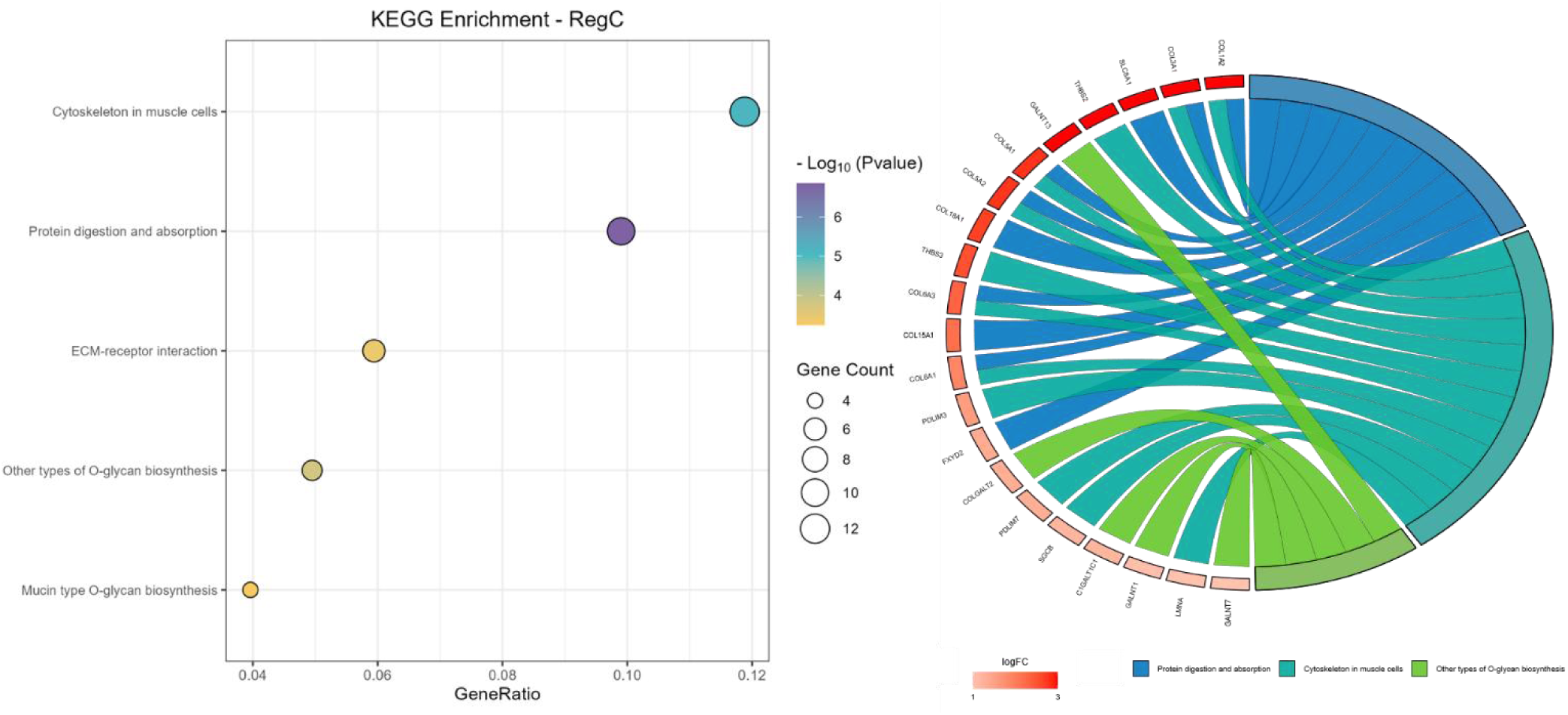
KEGG pathway enrichment of RegC OA-versus-control differential genes (FDR(q)<0.05). KEGG over-representation analysis results for RegC differential genes.

### Protein–protein interaction network analysis identifies collagen-centered hub genes

To prioritize candidate genes within the RegC OA-associated program, we constructed a protein–protein interaction (PPI) network using STRING and analyzed network topology in Cytoscape (Fig. 15). Degree-based ranking (cytoHubba) identified the top 5 hub genes as COL5A1, COL5A2, COL6A1, COL1A2, and COL3A1 (Fig. 15). These findings nominate a collagen-centered interaction core within the RegC OA-associated transcriptional program, complementing prior OA cartilage single-cell and multi-omics efforts that prioritize ECM-related modules and OA-critical populations for downstream validation.

**Figure 15.**
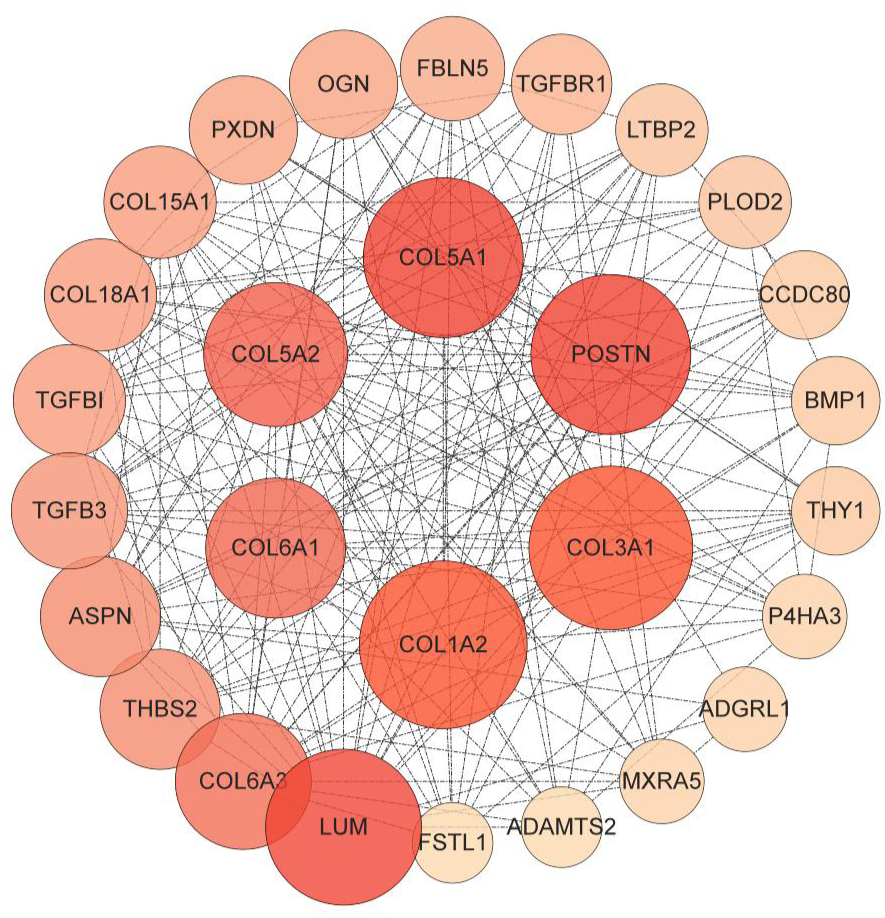
Protein–protein interaction network and hub gene prioritization in RegC. STRING-derived PPI network visualized in Cytoscape with degree-based hub ranking, highlighting top hub genes (COL5A1, COL5A2, COL6A1, COL1A2, COL3A1).

## Discussion

Osteoarthritis (OA) cartilage degeneration is increasingly recognized as a problem of cell-state remodeling rather than uniform failure of a single chondrocyte phenotype. In this context, single-cell transcriptomics has become central to defining disease-relevant chondrocyte populations and linking them to tractable molecular programs. Building on this framework, our reanalysis of the public human knee cartilage dataset GSE255460 delineated a chondrocyte state space with branching-like continuity and highlighted a regulatory chondrocyte (RegC) population positioned near the trajectory trunk and adjacent to multiple inferred branches. Within this transition-adjacent state, OA-versus-control differential expression converged on a coordinated program dominated by collagen-containing extracellular matrix (ECM) organization/remodeling, accompanied by endoplasmic reticulum (ER) lumen–associated secretory/proteostasis processes, cell–matrix adhesion (including focal adhesion), and transforming growth factor beta (TGF-β)/SMAD-related signaling. Network prioritization further nominated a collagen-centered hub module—COL5A1, COL5A2, COL6A1, COL1A2, and COL3A1—as high-connectivity candidates within the RegC OA-associated program. Taken together, these findings support the concept that OA transcriptional remodeling may be concentrated in transition-proximal chondrocyte states where matrix assembly, secretion/quality control, and adhesion–growth factor signaling are co-regulated, providing a focused set of candidate molecular nodes for downstream validation.

A key contribution of this study is the trajectory-informed prioritization of a transition-adjacent state, which provides a principled bridge between global chondrocyte heterogeneity and testable mechanistic hypotheses. Prior foundational work in human OA cartilage scRNA-seq described multiple chondrocyte populations and reported potential transitions among proliferative, prehypertrophic, and hypertrophic states, underscoring that OA involves coordinated state changes rather than isolated marker shifts^[4]^. More recently, multi-omics integration combining single-cell and spatial transcriptomics in human knee cartilage (including the same donor structure) defined an 11-population taxonomy with 33 population-specific markers and highlighted OA-critical populations such as inflammatory and prehypertrophic/hypertrophic states^[5]^. This broader single-cell literature consistently supports substantial chondrocyte heterogeneity and disease-critical subpopulations in OA cartilage^[1,12,13]^. Our reanalysis aligns with this trajectory-centric view of OA chondrocyte organization but specifically emphasizes that RegC is geometrically positioned at a convergence point in the inferred state manifold, adjacent to multiple branches. This “transition-adjacent” positioning does not establish lineage directionality; however, it provides a coherent rationale for focusing downstream differential and pathway analyses on a state that may be exposed to multiple remodeling trajectories within cartilage.

Within RegC, the OA-associated transcriptional program was most strongly characterized by collagen/ECM remodeling and organization, consistent with the central role of matrix disorganization in OA pathophysiology and with genetics and bulk transcriptomic studies that repeatedly implicate ECM pathways^[14,15]^. Large-scale OA GWAS integrating functional data reported enrichment for collagen formation and ECM organization pathways, reinforcing the relevance of collagen network remodeling as a core disease axis^[16]^. Independent functional-genomics resources in primary OA cartilage/chondrocytes further support genetically mediated regulation of cartilage programs relevant to OA risk and downstream pathways in disease tissue^[17–20]^. Bulk cartilage transcriptomic stratification also identified patient subgroups with differential activation of matrix-related and adhesion pathways, suggesting that ECM remodeling signatures are not merely end-stage epiphenomena but can define reproducible molecular phenotypes^[2,3]^. Our RegC-focused results extend these observations by locating collagen/ECM remodeling within a specific, transition-adjacent chondrocyte state, potentially sharpening the cellular context in which ECM regulation becomes coordinately rewired in OA.

Notably, the RegC OA-associated program included enrichment for ER lumen–associated secretory/proteostasis processes, which is biologically congruent with matrix remodeling as a high secretory-load state and with emerging OA literature emphasizing protein quality control as a modifiable disease mechanism^[21–23]^. A recent single-cell atlas of subchondral bone marrow lesions in OA highlighted proteostasis dysfunction and abnormal secretion of misfolded collagen as a druggable mechanism in early disease contexts (albeit in osteoblasts rather than chondrocytes), supporting the broader concept that proteostasis stress and matrix secretion can be mechanistically intertwined in OA tissues^[24]^. Additional single-cell osteochondral datasets further reinforce that cartilage degeneration is coupled to subchondral bone remodeling and bone–cartilage crosstalk, providing biological plausibility for convergent remodeling/secretory stress programs across joint compartments^[25–27]^. While our analysis does not assign causality, the co-enrichment of ECM remodeling and proteostasis/secretory pathways in RegC suggests a coherent module in which increased matrix synthesis/assembly demands may be accompanied by stress-handling programs that could shape the net balance between productive matrix deposition and maladaptive remodeling.

A further salient feature of the RegC signature was enrichment for cell–matrix adhesion/focal adhesion and TGF-β/SMAD-related signaling, pathways that are both mechanistically central and therapeutically nuanced in OA. TGF-β signaling has well-established context-dependent roles across the joint: dysregulated TGF-β activity in subchondral bone can initiate pathological remodeling and cartilage degeneration in vivo, demonstrating that the joint-wide TGF-β axis can be disease-driving under specific spatial and concentration contexts^[28]^. Within cartilage, multiple studies support TGF-β/SMAD signaling as a key regulator of matrix homeostasis and catabolic enzyme control. For example, TGFβ–SMAD2/3 signaling regulated FBXO6 expression to suppress MMP13 activation via MMP14 ubiquitination, directly connecting canonical signaling to matrix degradation control^[29]^. Conversely, cartilage matrix can regulate latent TGF-β sequestration and activation; CTGF served as a latent TGF-β binding protein controlling matrix sequestration and activation, with cartilage-specific deletion altering OA susceptibility^[30]^. Our finding that adhesion-related terms and TGF-β/SMAD-related signaling co-enrich with ECM remodeling in RegC is therefore consistent with a model in which matrix assembly, mechanotransduction/adhesion, and growth factor signaling are coordinated within transition-proximal chondrocyte programs.

Network analysis provided a practical prioritization step by nominating a collagen-centered hub module (COL5A1, COL5A2, COL6A1, COL1A2, COL3A1). Importantly, hub status reflects high connectivity in the inferred interaction network, not proven causality. Nevertheless, the specific identities of these hubs are biologically plausible in OA given accumulating evidence implicating collagen network architecture and pericellular matrix composition in chondrocyte mechanobiology and disease states. A recent mechanistic organoid study demonstrated that a damaging COL6A3 variant altered pericellular matrix interactions and provoked an osteoarthritic chondrocyte state, illustrating how collagen VI–related perturbations can shift chondrocyte phenotypes under mechanical stress contexts^[31]^. In parallel, OA multi-joint spatiotemporal multi-omics identified a COL5A1+ fibroblast population contributing to fibrocartilaginous ECM transformation and implicated integrin activity as a regulator of aberrant connective tissue remodeling, supporting the broader relevance of COL5A1-associated matrix programs in OA-related tissue remodeling^[32]^. These external lines of evidence do not directly validate our RegC hubs, but they strengthen the rationale that collagen-network nodes prioritized from a transition-adjacent chondrocyte program may be meaningful candidates for targeted validation (e.g., protein-level localization, spatial co-expression with RegC markers, and perturbation studies).

Our findings should also be interpreted in the broader landscape of OA single-cell studies that increasingly identify disease-relevant subpopulations with distinct regulatory logic. For instance, a senescent pathogenic subset shared between cartilage and meniscus was expanded in OA and acted as a dominant signaling sender/receiver, including reception of TGFβ signaling, with transcriptional regulators such as ZEB1 implicated as key drivers^[13]^. Orthogonal single-cell proteomic profiling of OA cartilage revealed rare inflammation-amplifying and inflammation-dampening subpopulations and proposed pharmacologic targeting strategies, emphasizing that actionable biology can be concentrated in specific, sometimes rare, cellular states^[12]^. In this setting, our RegC-centered results add a complementary dimension: rather than focusing on inflammatory amplification alone, we highlight a transition-adjacent remodeling program in which ECM/collagen organization is central and is coupled to proteostasis and adhesion–TGF-β signaling, potentially representing a mechanistically coherent “remodeling junction” in the chondrocyte state space.

Several limitations temper interpretation and define the most appropriate next validation steps. First, this study is a single-dataset reanalysis of a public cohort; while internal consistency across multiple analytic layers (trajectory position → differential expression → enrichment → network hubs) strengthens plausibility, external replication in independent human cartilage scRNA-seq datasets would be essential to establish generalizability. Second, trajectory inference methods (including Slingshot) recover state adjacency and branching structure but do not, on their own, establish true lineage directionality or causal transitions; thus, RegC should be interpreted as transition-adjacent rather than definitively “transitional” in a developmental sense. Third, PPI-derived hub prioritization is inherently model- and database-dependent; hub genes should be treated as candidates for validation rather than definitive drivers. Fourth, the current analysis centers on transcriptomic readouts; protein-level and spatial validation within cartilage zones would be particularly informative, given prior evidence that OA-associated differential genes can be spatially concentrated in specific cartilage regions.

## Conclusion

In summary, this reanalysis supports a model in which OA-associated transcriptional remodeling is concentrated within a transition-adjacent RegC state that coordinates collagen/ECM organization, secretory/proteostasis stress programs, adhesion, and TGF-β/SMAD-related signaling, and it nominates a collagen-centered hub module (COL5A1/COL5A2/COL6A1/COL1A2/COL3A1) for downstream validation. By coupling consistent state annotation with trajectory-informed prioritization, the study offers a focused and testable set of hypotheses for how matrix remodeling programs may be organized in transition-proximal chondrocyte states in OA.

## Declarations

Ethics and Consent to Participate declarations: not applicable. This study is a retrospective re-analysis of a publicly available human single-cell RNA-sequencing dataset (GSE255460). The original data were collected from human knee cartilage samples with proper ethical approval and informed consent, as reported in the original publication. Our study did not involve any direct contact with human subjects, recruitment of participants, or collection of new biological samples. Therefore, additional ethics approval and consent to participate are not applicable for this work.

